# A helicase-containing module defines a family of pCD630-like plasmids in *Clostridium difficile*

**DOI:** 10.1101/198473

**Authors:** Wiep Klaas Smits, J. Scott Weese, Adam P. Roberts, Céline Harmanus, Bastian Hornung

## Abstract

*Clostridium difficile* is a Gram-positive and sporulating enteropathogen that is a major cause of healthcare-associated infections. Even though a large number of genomes of this species have been sequenced, only a few plasmids have been described in the literature. Here, we use a combination of *in silico* analyses and laboratory experiments to show that plasmids are common in *C. difficile*. We focus on a group of plasmids that share similarity with the plasmid pCD630, from the reference strain 630. The family of pCD630-like plasmids is defined by the presence of a conserved putative helicase that is likely part of the plasmid replicon. This replicon is compatible with at least some other *C. difficile* replicons, as strains can carry pCD630-like plasmids in addition to other plasmids. We find two distinct sub-groups of pCD630-like plasmids that differ in size and accessory modules. This study is the first to describe a family of plasmids in *C. difficile*.

## Introduction

*Clostridium difficile* (*Clostridioides difficile* [1]) is a Gram-positive, anaerobic and spore-forming bacterium that can asymptomatically colonize the human gut [2]. It is ubiquitous in the environment, and can also be found in the gastrointestinal tract of many animals. The bacterium gained notoriety when it was identified as the causative agent of health care associated diarrhea, and is increasingly implicated in community-associated disease in many countries [2]. In hosts with a dysbiosis of the microbiome, such as patients treated with broad-spectrum antimicrobials, conditions are favorable for *C. difficile* germination and outgrowth [3]. *C. difficile* produces one or more toxins, that cause symptoms ranging from diarrhea to potentially fatal toxic megacolon [2, 4].

Over the past two decades, genetic studies have of *C. difficile* have become possible due to the generation of shuttle plasmids that can be transferred by conjugation from *Escherichia coli* to *C. difficile* [5]. These plasmids mostly employ a replicon derived from plasmid pCD6 for replication in *C. difficile* [6]. In 2006, the first genome sequence of *C. difficile* became available, revealing the presence of another plasmid, pCD630 [7].

Despite a great number of strains having been whole genome sequenced since then, plasmid biology of *C. difficile* has been poorly explored. One reason is that plasmid content is variable, and most studies on the evolution and/or transmission of *C. difficile* focus on those genes conserved between all strains (the core genome) [8–11]. However, there is reason to assume that plasmids are common in *C. difficile*; for instance, before the advent of the currently common typing schemes [12], plasmid isolation had been proposed as an epidemiological tool [13]. The ratio of plasmid-containing to plasmid-free strains in this study was found to be approximately 1:2, suggesting that around 30% of strains of *C. difficile* may carry a plasmid. Furthermore, hybridization-based analyses of total DNA from a collection of *C. difficile* strains suggest the presence of DNA with significant similarity to pCD630 open reading frames (ORFs) [14, 15].

Here, we define a family of plasmids that share a conserved helicase-containing module and demonstrate that these plasmids are common in a diverse set of *C. difficile* strains.

## Materials and methods

### Strains and growth conditions

Strains used in this study are listed in Table 1. For DNA isolation, strains were grown on *Clostridium difficile* agar (CLO) plates (bioMérieux) and a single colony was inoculated into pre-reduced brain-heart infusion broth (Oxoid) supplemented with 0.5 % w/v yeast extract (Sigma-Aldrich) and *Clostridium difficile* selective supplement (Oxoid). Strains were PCR ribotyped [16] in-house for this study. For some strains a PCR ribotype could not be assigned (Table 1).

**Table 1.**
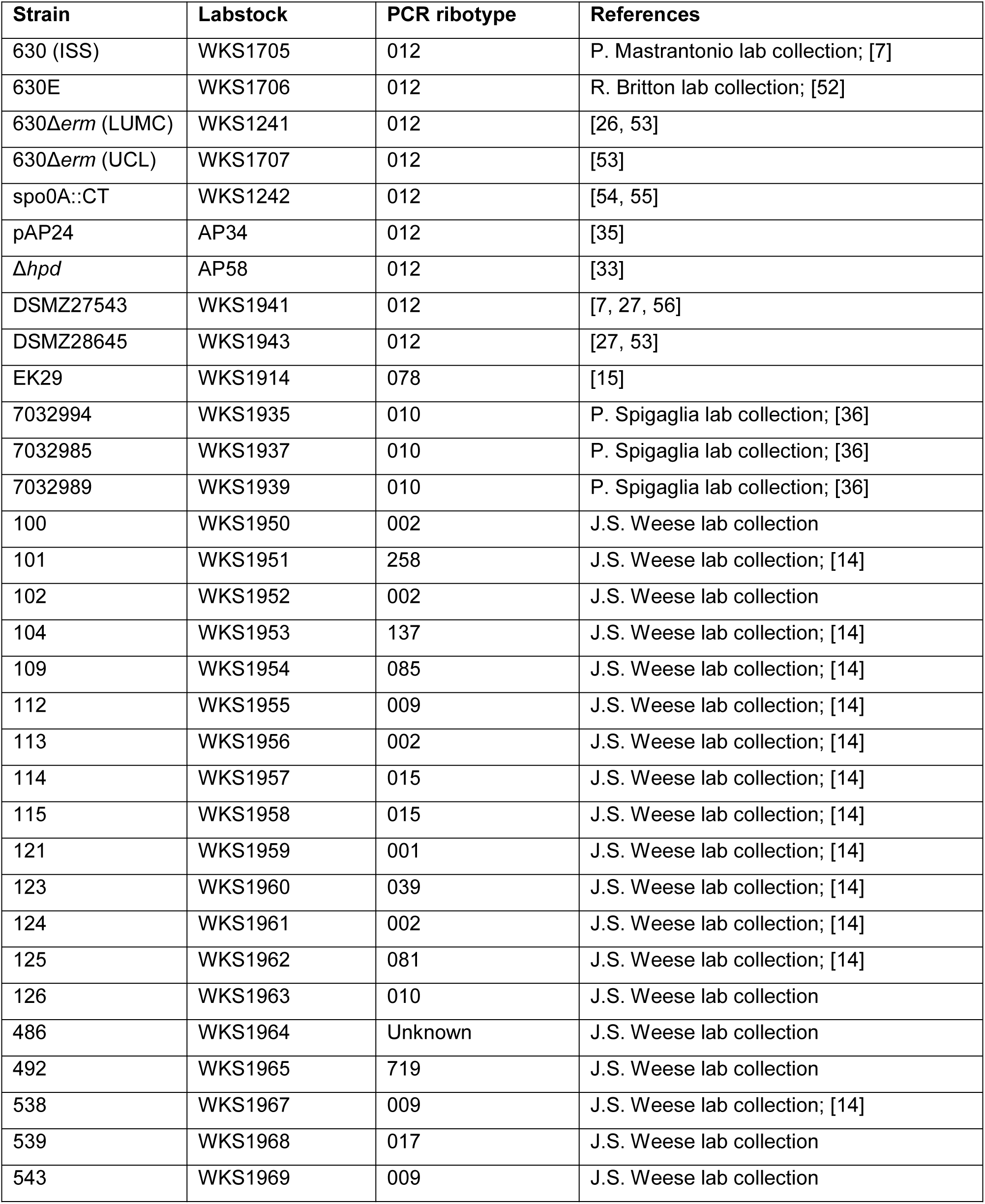
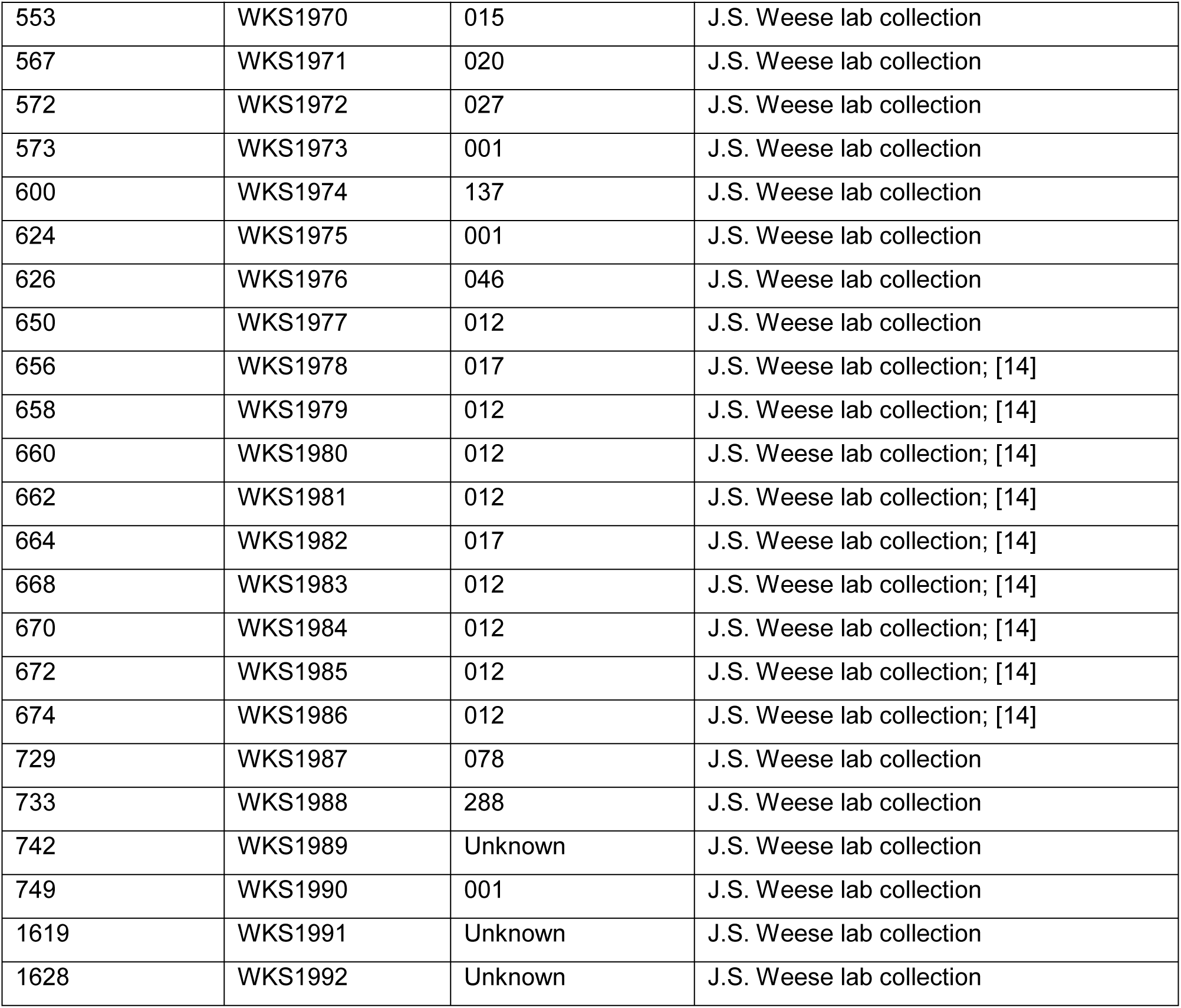
*S*trains used in this study.

### Isolation of plasmid DNA from C. difficile

Plasmids were isolated from 2 mL of *C. difficile* overnight culture using NucleoSpin Plasmid Easypure columns (Macherey-Nagel); to increase yield, 10mg/mL lysozyme to buffer A1 was added, as recommended by the manufacturer. Using PCR and sequencing, we found that the DNA isolated using this kit is heavily contaminated with chromosomal DNA. To isolate pure plasmid DNA, aliquots of the DNA were incubated with PlasmidSafe ATP-dependent DNase (Epicentre) that digests linear, but not circular double stranded DNA. After purification with a Nucleospin Gel and PCR Clean-up kit (Macherey-Nagel), the absence of genomic DNA was confirmed by PCR using primers targeting *gluD* (Table 2). Yields were generally very low, but the plasmid was readily detectable by PCR.

**Table 2.**
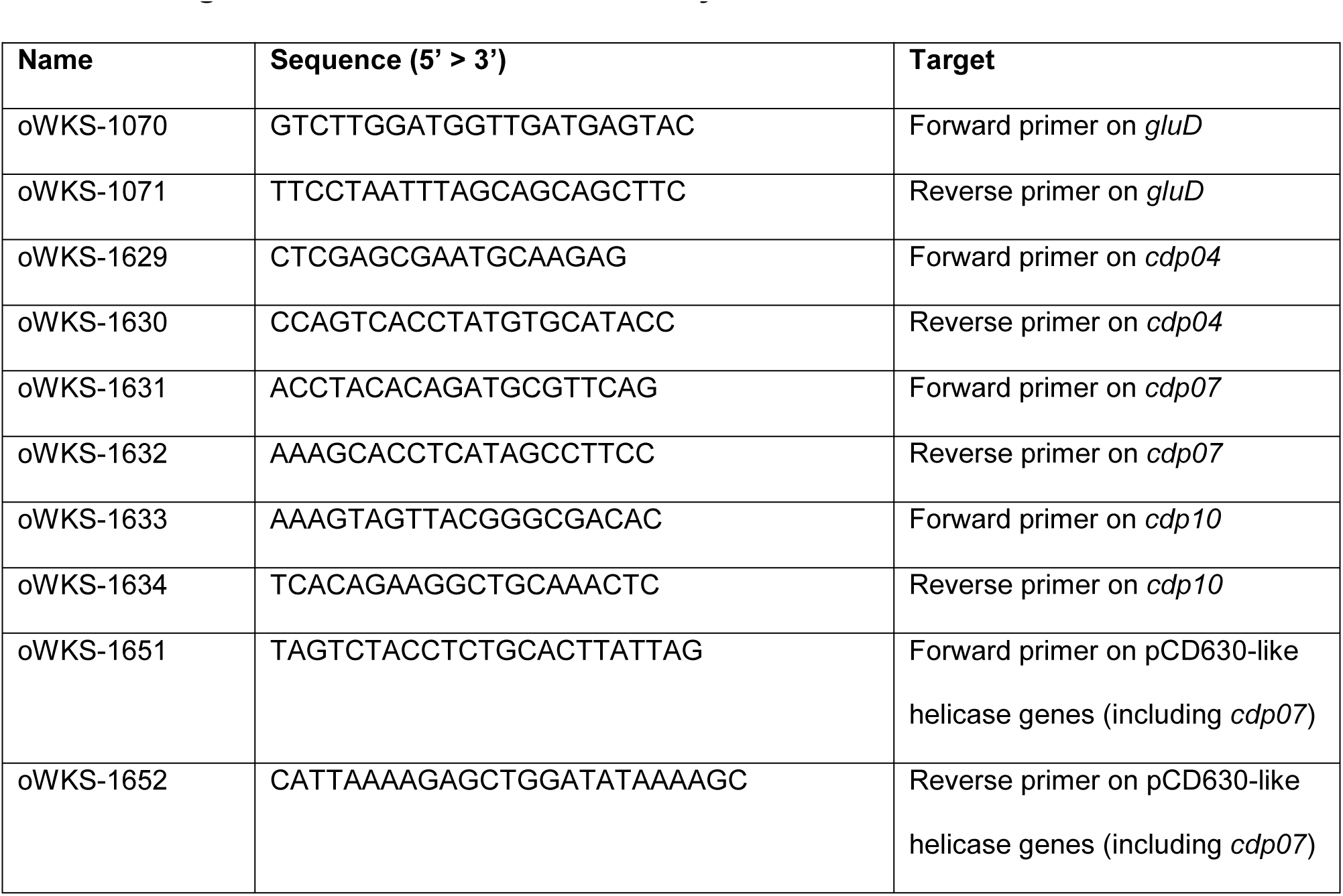
Oligonucleotides used in this study.

### Reannotation of pCD630 and identification of a pCD630-like plasmid

The pCD630 sequence was obtained from GenBank (AM180356.1). CDP01 and CDP11 form a single open reading frame (ORF) and were treated as a single ORF in our analyses. Protein sequences encoded by the ORFs of pCD630 were used as BLAST queries against the non-redundant protein sequences database, limited to taxid:1496 (*Clostridium difficile*). This identified the 8089 bp *Peptoclostridium difficile* genome assembly 7032985, scaffold BN1096_Contig_85 (LK932541.1). To reconstitute the plasmid from this contig, the DNA was circularized and a single copy of the 98 bp direct repeat that was present at the terminus of the original contig was removed using Geneious R10. The resulting 7991 bp sequence now encodes a full copy of a sequence homologous to CDP07 of pCD630. Reannotation of plasmids was performed using using an in-house pipeline. This pipeline incorporates the gene caller Prodigal (version 2.6.3) [17], RNAmmer (version 1.2) [18], Aragorn (version 1.2.38) [19], the CRISPR recognition tool (version 1.2) [20], dbCAN (version 5.0) [21] and PRIAM (version March 2015) [22]. The plasmid derived from LK932541 was submitted to GenBank as pCD-ISS1 (GenBank: MG266000).

### Identification of pCD630-like plasmids in short read archives

In order to identify other pCD630-like plasmids in sequence databases, paired end Illumina sequences from study PRJEB2101 (ERR017367-ERR017371, ERR022513, ERR125908-ERR125911) were downloaded from the short read archive of the European Nucleotide Archive (ENA). Short reads were assembled and visualized in PLACNETw [23] to determine likely replicons. The contigs corresponding to pCD630-like plasmids were downloaded and imported into Geneious R10 software (Biomatters Ltd) for circularization and removal of terminal repeats; afterwards all plasmids which could be circularized were compared with BLASTN (version 2.40) [24] to pCD630 and the sequences were restructured to start at the base corresponding to base 2903 in pCD630. Afterwards the plasmids were annotated using the in-house pipeline as described above, and submitted to GenBank as pCD-WTSI1 (GenBank: MG019959), pCD-WTSI2 (GenBank: MG019960), pCD-WTSI3 (GenBank: MG019961), pCD-WTSI4 (GenBank: MG019962). Alignments of the pCD630-like plasmids were performed in Geneious R10 (Biomatters Ltd) and the alignment figure was prepared using Adobe Illustrator CC (Adobe Systems Inc).

### Polymerase chain reaction

Oligonucleotides used in this work are listed in Table 2. To confirm the presence of pCD630 in derivatives of *C. difficile* strain 630, PCR was performed with oWKS-1629 and oWKS-1630 (targeting CDP04); oWKS-1631 and oWKS-1632 (targeting CDP07); oWKS-1633 and oWKS-1634 (targeting CDP10). As a control for chromosomal DNA, a PCR was performed targeting the *gluD* gene that is used as a target for *C. difficile* identification, using primers oWKS-1070 and oWKS-1071. To screen a collection of *C. difficile* strains for the presence of pCD630-like plasmids, a PCR was performed with primers oWKS-1651 and oWKS-1652 that targets a region of CDP07 conserved among the 6 full length plasmids identified in this work. Fragments were separated on 0.5x TAE (20 mM Tris, 10mM acetic acid, 0.5mM EDTA) agarose, stained with ethidium bromide and imaged on a Gel Doc XR system (BioRad). Images were captured using QuantityOne (BioRad) and prepared for publication using Adobe Photoshop CC (Adobe Systems Inc) and CorelDRAW X8 (Corel Corporation).

## Results and discussion

### pCD630 is present in C. difficile strain 630 and some of its derivatives

The most commonly used laboratory strains of *C. difficile* are derived from the reference strain 630 (PCR ribotype 012 [7]) by serial passaging and screening for loss of resistance to the antimicrobial erythromycin [25]). It was generally assumed that during this passaging the plasmid pCD630 that is present in strain 630 was lost. Indeed, our *de novo* assembly of the 630Δ*erm* genome sequence using single molecule real-time (SMRT) sequencing did not report the presence of the plasmid [26]. Recently, however, one study showed the presence of reads mapping to pCD630 in a genome resequencing project of another isolate of 630Δ*erm* [27]. This prompted us to revisit our whole genome sequencing data (ENA:PRJEB7326). If the plasmid was maintained in 630Δ*erm*, we expected to be able to find reads mapping back to the pCD630 reference sequence (GenBank: AM180356.1) in this dataset. Indeed, when we performed a reference assembly of the short reads (ENA: ERR609091) against the pCD630, we found approximately 0.8% of the reads mapping to the plasmid. The original *de novo* assembly overlooked the plasmid due to a low number of plasmid-mapping reads as the result of a size fractionation step (the plasmid is <8kb, and SMRT sequencing was performed on high MW DNA). Notably, both a *de novo* assembly of the plasmid based on a small number of SMRT reads, as well as the reference assembly using a large number of Illumina reads shows a 100% congruence with the published reference sequence for pCD630 (data not shown). This indicates that, despite the lack of selective pressure and repeated culturing under laboratory conditions, the plasmid has remained unchanged.

We confirmed the presence of pCD630 and the extrachromosomal nature of the plasmid. To do so, we performed a miniprep on a *C. difficile* liquid culture and treated the resulting DNA with PlasmidSafe DNase, that selectively removes linear double stranded (sheared) but not circular DNA. A PCR using primers against three ORFs of pCD630 (*cdp04, cdp07 and cdp10*) and one chromosomal locus (*gluD*) showed that the DNase treated samples were negative for the *gluD* PCR, but positive for all three plasmid loci (Figure 1A).

**Figure 1.**
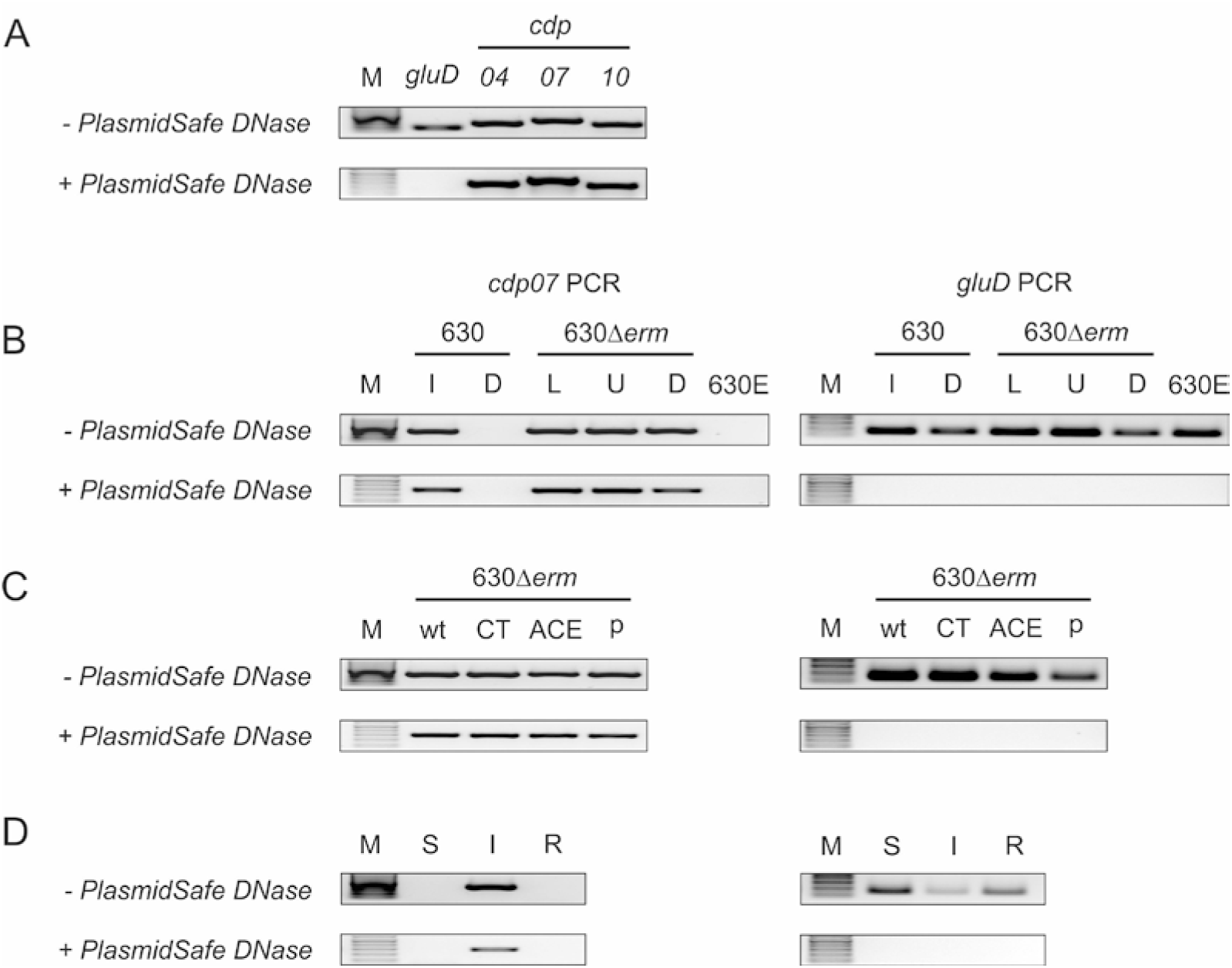
630 and derivatives can contain pCD630. **A.** *C. difficile* 630Δ*erm* [26] contains the pCD630 plasmid. **B.** Some, but not all, 630-derived strains contain pCD630. I=ISS D=DSMZ L=LUMC U=UCL [26]. **C.** Genetically modified 630Δ*erm* strains still contain pCD630. wt = wild type, CT = Clostron mutant [30, 31], ACE = allelic coupled exchange mutant [32, 33], p = containing a replicative plasmid [34, 35]. **D.** Strain 7032985 (intermediate metronidazole susceptible; I) contains a pCD630-like plasmid but strains 7032994 (metronidazole susceptible; S) and 7032989 (metronidazole resistant; R) do not. For oligonucleotides used, see Materials and Methods. M = marker.

The results above suggest that pCD630 is stably maintained extrachromosomally. Next, we wanted to verify the presence of the plasmid in multiple derivatives of strain 630, to see if plasmid-loss could be documented. We previously analyzed 630Δ*erm* from our laboratory as well as from the laboratory where it was generated to determine the chromosomal location of the mobile element CTn*5*, in comparison with the ancestral strain 630 and the independently derived 630E strain [26]. We found that pCD630 was readily detected on total genomic DNA from all these strains, with the exception of the 630E isolate in our collection (Figure 1B). 630E and 630Δ*erm* demonstrate notable phenotypic differences [25, 28] and we wondered whether these might be in part due to loss of the pCD630 plasmid. We performed a reference assembly using the whole genome sequencing data available from the study by Collery *et al* (ENA: PRJNA304508), that compares 630Δ*erm* and 630E [28]. The assembly showed that both these strains contain pCD630 and indicate that the loss of plasmid is not a general feature of 630E strains. We conclude that the observed phenotypic differences are not likely due to loss of the plasmid. It was reported that the isolate of *C. difficile* 630 stored at in the collection of the DSMZ (www.dsmz.de) lacks the pCD630 plasmid [27, 29]. We requested both 630 (DSMZ 26845) and 630Δ*erm* (DSMZ 27543) and checked for the presence of the plasmid. Our results confirm the absence of pCD630 from DSMZ 26485 (Figure 1B), in line with the analysis of Dannheim *et al* [27].

In other organisms, the presence of certain replicons can negatively affect the maintenance of other replicons (plasmid incompatibility); this has not been documented for *C. difficile* to date. If pCD630 would be incompatible with other replicons (such as the pCB102 and pCD6) [5, 30], this could result in loss of the pCD630 plasmid in genetically modified *C. difficile*. We therefore tested whether pCD630 was lost in strains chromosomally modified using Clostron mutagenesis [30, 31], Allele Coupled Exchange [32, 33] or carrying a replicative plasmid [34, 35]. We found that all of these carried pCD630, suggesting that pCD630 is compatible with pCB102 and pCD6-based replicons (Figure 1C). Similar results were obtained with multiple mutants (data not shown).

Together, our data clearly shows that pCD630 persists in the absence of selection, but also that pCD630 <u>can</u> be lost. Thus, care should be taken to verify plasmid content when comparing presumed isogenic laboratory strains even when they are derived from the same isolate.

### A pCD630-like plasmid is present in a strain with reduced metronidazole susceptibility

We wondered whether there are more pCD630-like plasmids. As a first step, we set out to identify coding sequences with homology to pCD630 ORFs in GenBank. Using default settings, we identified a single 8089 bp contig that encodes proteins with homology to CDP01, CDP04-6 and CDP08-11 (*Peptoclostridium difficile* genome assembly 7032985, scaffold BN1096_Contig_85; GenBank: LK932541) (Figure 2).

**Figure 2.**
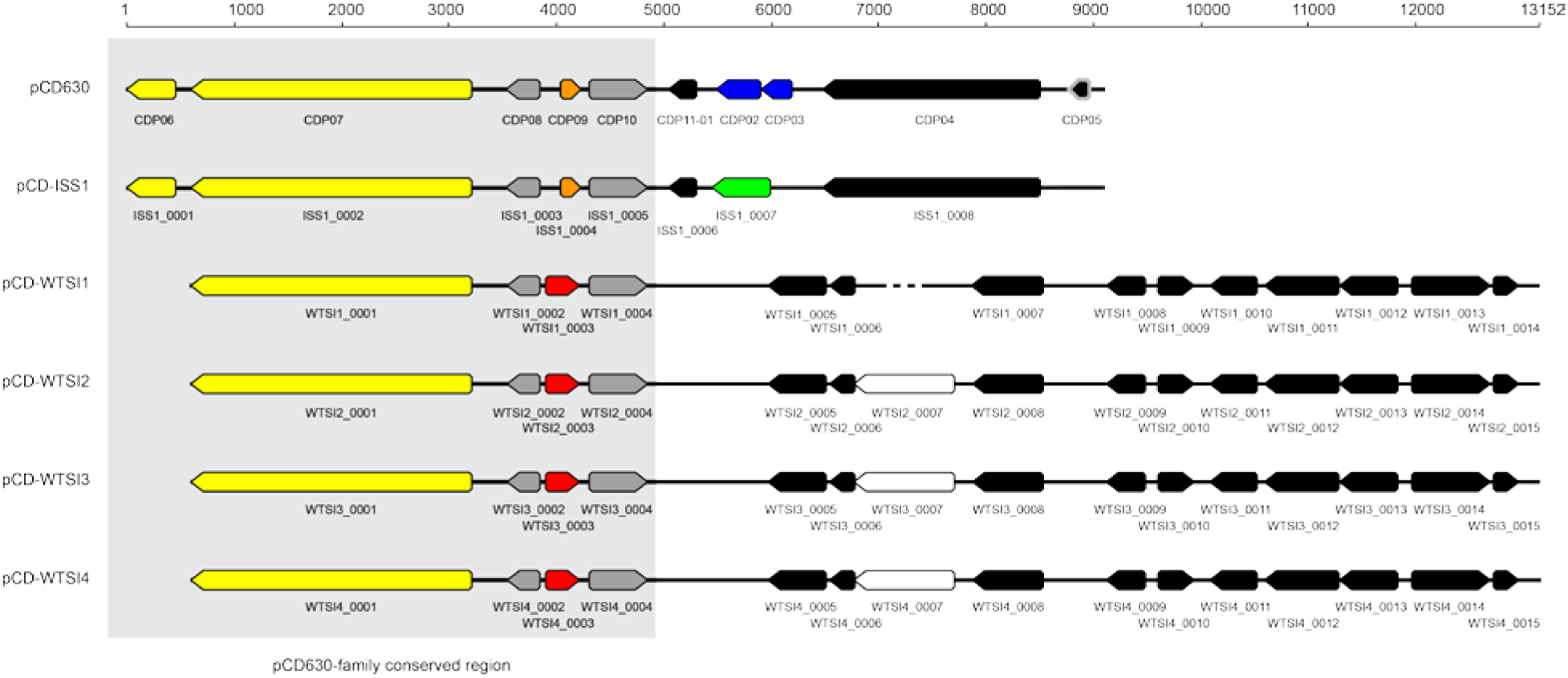
Schematic representation of an alignment of pCD630-like plasmids. Full-length plasmids identified in this study were aligned. pCD-ISS1 is based on GenBank:LK932541. pCD-WTSI-1 to pCD-WTSI4 are based on short read sequences from ENA:PRJEB2101. The most striking differences are indicated with differently colored ORFs. The conserved module encompassing the gene encoding a helicase is boxed, the accessory module is indicated with black ORFs. The gray outline of CDP05 indicates it is annotated in AM180356.1 but is not predicted in our analysis.

This sequence stems from a study that compares three non-toxigenic PCR ribotype 010 strains of *C. difficile*, with differing susceptibility to metronidazole [36]. Strain 7032985 was classified as intermediate resistant to metronidazole. If we assume that the contig represents a pCD630-like plasmid, we expect DNA from this strain to remain positive in a PCR that targets the plasmid upon treatment with PlasmidSafe DNase. We found that the PCR targeting *cdp07*, but not chromosomal locus *gluD,* results in a clear signal when using a template treated with PlasmidSafe DNase (Figure 1D). Having confirmed that the contig is extrachromosomal in nature, we will refer to the putative plasmid as pCD-ISS1 hereafter (Table 3).

**Table 3.** Full length pCD630-like plasmids.

| Name | Source | Accessions | Size | Reference |
| --- | --- | --- | --- | --- |
| pCD630 | Strain 630 | GenBank: AM180356 | 7881 bp | [7]; this study |
| pCD-ISS1 | Strain 7032985 | GenBank: LK932541 (contig)<br>GenBank: MG266000 (plasmid) | 7991 bp | [36]; this study |
| pCD-WTSI1 | Not specified | ENA: ERR017368 (Illumina reads)<br>GenBank: MG019959 | 11777 bp | This study |
| pCD-WTSI2 | Not specified | ENA:ERR022513 (Illumina reads)<br>GenBank: MG019960 | 12526 bp | This study |
| pCD-WTSI3 | Not specified | ENA: ERR125910 (Illumina reads)<br>GenBank: MG019961 | 12525 bp | This study |
| pCD-WTSI4 | Not specified | ENA: ERR125911 (Illumina reads)<br>GenBank: MG019962 | 12488 bp | This study |

To further analyze pCD-ISS1, we circularized the LK932451 contig to yield a putative plasmid of 7991bp, performed an automated annotation (GenBank: MG266000) and compared the annotated pCD-ISS1 sequence to that of pCD630 (Figure 2).Overall, the two plasmids are highly similar. Of note, the ORF that corresponds to the DEAD/DEAH helicase like protein (CDP07 in pCD630) was not annotated in the LK932541 contig due to its linear nature, but is evident in the pCD-ISS1 sequence. Similarly, we found that CDP1 (gene remnant) and CDP11 (doubtful CDS) of pCD630 are in fact a single 201bp ORF, as annotated in the LK932541 contig. A revised annotation of pCD630 has been submitted to ENA (AM180356) to reflect this. Though the pCD-ISS1 and pCD630 plasmids are co-linear, there is a single region that is divergent. The region of pCD630 encompassing the ORFs encoding CDP02 and CDP03 is absent from pCD-ISS1; the latter contains an ORF encoding a RNA polymerase sigma factor protein (Interpro:IPR013324) in this region. The pCD-ISS1 annotation does not identify an ORF encoding a homolog of CDP05 of pCD630. This is the result of a 2bp deletion; it suggests that CDP05 (previously annotated as a doubtful CDS) may not be a true coding sequence. Both pCD630 and pCD-ISS1 encode phage-related functions. Most notably, CDP04 and its homolog encode a phage capsid protein with similarity to the HK97-like major capsid proteins of tailed phages of the Caudovirales order. Caudovirales are common *C. difficile* phages [37–40]. However, beside the phage capsid, pCD630 and pCD-ISS11 lack genes encoding other proteins required for virion formation, such as the large terminase subunit and the portal protein. Therefore, it is highly unlikely that phage particles can be produced from these plasmids. In line with this, we find that the genes encoding the phage proteins are poorly, if at all, expressed (unpublished observations). It appears therefore that (part of) a viral genome has been incorporated into the plasmid, or that the viral genome has been transformed into a plasmid.

Together, these data suggest the existence of plasmids closely related to pCD630 in at least two different PCR ribotypes (010 and 012).

### pCD630-like plasmids can be identified in short reads from whole genome sequence projects

Above, we showed the existence of at least one pCD630-like plasmid. We wondered if we could extend the family by interrogating the wealth of raw, non-annotated, sequence data in the public domain. We downloaded a selection of sequence reads from ENA, corresponding to 10 different strains (see Materials and Methods). To identify extrachromosomal replicons, we used graph-based tool for reconstruction of plasmids [23]. We validated this tool on our short read sequence data from our 630Δ*erm* sequence (ERR609091)[26] and found that is readily identifies the pCD630 plasmid (data not shown).

Surprisingly, we found only two strains with a single replicon (i.e. only the chromosome). The other 8 analyzed datasets suggested the presence of at least one other replicon. Strikingly, 6 contained a replicon that shared similarity to pCD630. Of these, 4 could be circularized due to the presence of direct repeats at the ends of the contig and therefore likely represent complete plasmid sequences, as was the case for pCD-ISS1 (Table 3). These plasmids - hereafter referred to as pCD-WTSI1, pCD-WTSI2, pCD-WTSI3 and pCD-WTSI4 – are all significantly larger than pCD630 and pCD-ISS1 (Figure 2). The smaller pCD630-like contigs without flanking repeats (that may represent either complete, or incomplete plasmids) were not further studied.

To gain further insight in the group of large pCD630-like plasmids, we performed an automated annotation of plasmids pCD-WTSI1 (GenBank: MG019959), pCD-WTSI2 (GenBank: MG019960), pCD-WTSI3 (GenBank: MG019961) and pCD-WTSI4 (GenBank: MG019962). The homology with the small pCD630-like plasmids is confined to the region encoding CDP6-CDP10 of pCD630. Within this region, it is noteworthy that the ORF encoding the Arc-type ribbon-helix-helix protein (Pfam: PF12651) CDP09 of pCD630 appears to be replaced with another putative DNA binding protein, a helix-turn-helix XRE protein (InterPro:IPR010982) in the pCD-WTSI group of plasmids. Further, we noted that the CDP06, that encodes a truncated homolog of CDP07, appears to be fused with CDP07 to form a hybrid protein nearly identical in size to CDP07. This suggests that the CDP06-07 arrangement may be the result of an (incomplete) gene duplication event. The proteins are putative superfamily 2 helicase fused to an N-terminal CHC2 zinc finger domain, with homology to the corresponding TOPRIM domain of DnaG-like primases. They also contains a third domain of unknown function C-terminal of the helicase domain.

The pCD-WTSI plasmids all contain a highly similar accessory module of ~8kb. Within this module, notable functions include an integrase (Pfam: PF00589), a recombinase (Pfam: PF00239), a Cro-C1-type HTH protein (Pfam: PF01381), a penicillinase repressor (Pfam: PF03965), and an RNA polymerase sigma factor (Pfam: PF08281 & Pfam: PF04542). The combination is suggestive of integration of mobile genetic element(s) into the plasmid backbone.

In the short read archive, we only identified large pCD630-like plasmids so far.

Though we cannot exclude the existence of more small pCD630-like plasmids, we consider it likely that the pCD-WTSI plasmids represent a more widely distributed form of the pCD630-like plasmid family.

### pCD630-like plasmids have a modular organization

Above, we have identified 6 plasmids sharing significant homology in a region that encompasses an ORF encoding a putative helicase. Moreover, we have shown that the large and small pCD630-like plasmids are remarkably similar, but that certain genes appear to have been exchanged. Thus, the organization of these plasmids, like those of mobile elements in *C. difficile* [41, 42] and plasmids in other organisms [43], is modular.

None of the pCD630-like plasmids encodes a previously characterized replication protein; yet, it is clear that the plasmid is efficiently maintained in the absence of obvious selection (Figure 1). Based on the finding that all 6 plasmids contain homologs of the pCD630 CDP6-10, we propose that this region (or part of it) forms the replicon of the plasmids. The DEAD-DEAH family helicase CDP07 and its homologs, that also contain a CHC2 zinc finger domain (InterPro: IPR002694) that aligns with the corresponding domain in DnaG-like DNA primases, appear to be the most likely candidate to be the replication proteins for this family of plasmids. As noted above, in the large pCD630-like plasmids the helicase is a CDP06-7 hybrid protein; this may underlie the signals corresponding to these ORFs in microarray and comparative genome hybridization studies, but also suggests that CDP06 itself is probably dispensable for plasmid maintenance. CDP09 is likely also not crucial for the function of the replicon, as it is replaced by another protein in the group of large pCD630-like proteins. It is conceivable that CDP09 and the HTH XTRE proteins serve a regulatory function for instance in controlling the copy number of the plasmids. The small pCD630-like plasmids have an estimated copy number of 4-5, based on average read coverage for chromosomal loci and the plasmid contigs. For the large plasmids, this is 9-10. Consistent with a regulatory rather than an essential function, we noted that in a previous microarray identification more strains appear to contain homologs of CDP6-10 than any of the other pCD630 genes, and that several strains harboring CDP6-8 and CDP10 do not contain CDP09 [14]. The same study also found strains that carry homologs of CDP02-03, but not any of the other genes of pCD630. Combined with our observation that this particular region is replaced with a single ORF in pCD-ISS1, suggest that CDP02-03 have been horizontally acquired. In line with this notion, we found that CDP02 has homology to HNH endonucleases (PFAM01844.17), and genes encoding these homing endonucleases are considered as selfish genetic elements [44].

### pCD630-like plasmids are common in diverse ribotypes

The identification of 6 plasmids carrying a conserved putative replication module, allowed us to determine the most conserved regions within this module. We designed primers against one such region, to be able to identify pCD630-like plasmids by PCR. We tested these primers in a PCR reaction on chromosomal DNA from strains 630Δ*erm* (WKS1241), yielding a positive signal (Figure 3). Next, we tested a collection of 43 strains of diverse PCR ribotypes to see if pCD630-like plasmids could be identified. We found DNA from 11 isolates gave a signal similar or greater than our positive control, 630Δ*erm* (32.6%); this includes strains of PCR ribotypes 012, 015, 017, and 081 (Figure 3). Interestingly, strain 630 and derivatives are PCR ribotype 012 as well [7].

**Figure 3.**
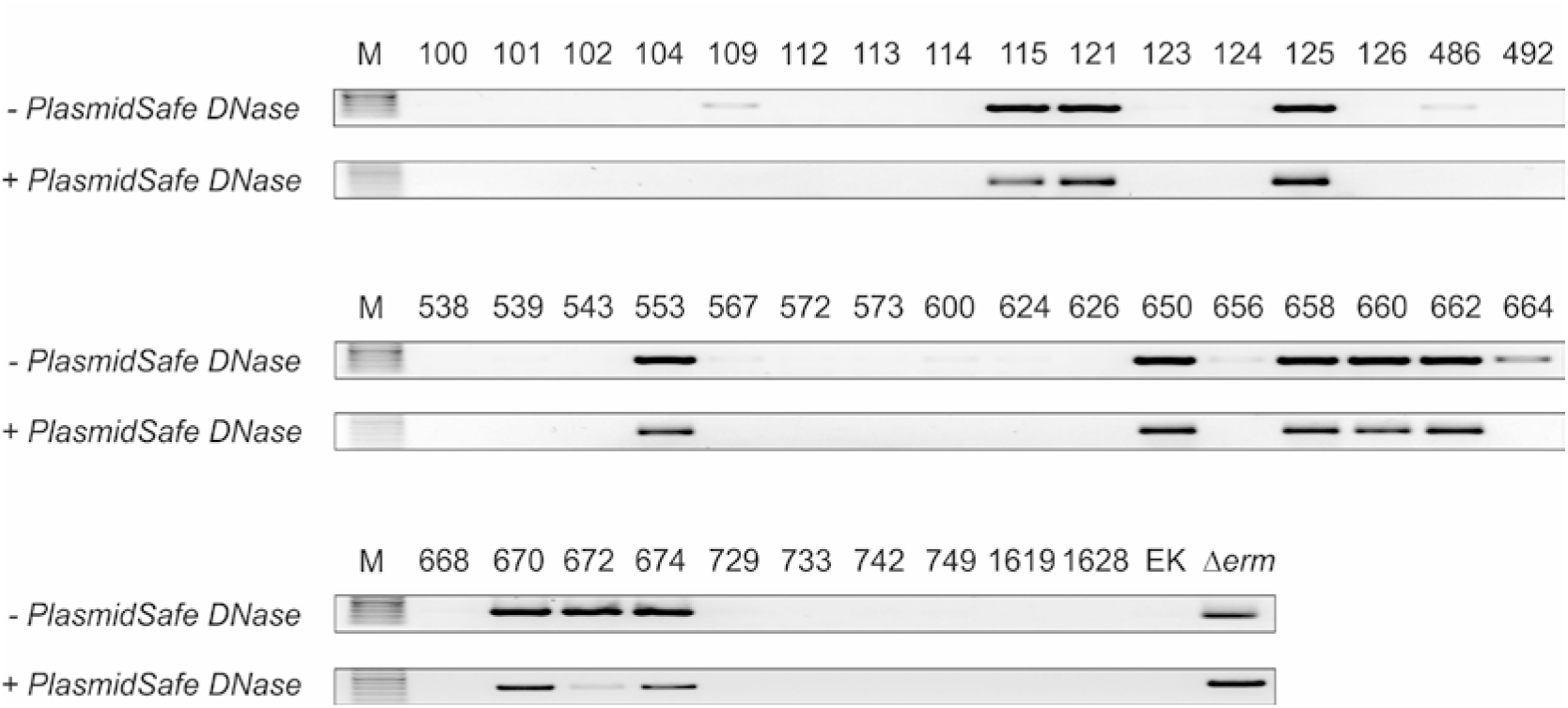
pCD630-like plasmids are present in diverse *C. difficile* strains. A PCR was performed against a conserved target region in the putative helicase protein using oWKS-1651 and oWKS-1652. The presence of a pCD630-like plasmid results in a positive signal in this PCR. M = marker, EK = EK29 [15], Δ*erm* = 630Δ*erm* [26].

Those samples that were weakly positive on total DNA, appear negative on PlasmidSafe DNase treated DNA and are therefore likely false positives. Alternatively, these could represent isolates in which the plasmid is integrated into the chromosome. Isolating and characterizing these plasmids is part of our ongoing work. We noted that strain EK29, that presumably contains a pCD630-like plasmid [15], appears negative in this PCR. We interpret this to mean that the PCR likely fails to detect certain pCD630-like plasmids, suggesting that the actual number of strain containing pCD630-like plasmids may be even higher. Our data suggests that pCD630-like plasmids are common, and not limited to PCR ribotype 010 (strain 7032985) and 012 (strains 630 and derivatives).

The high prevalence of pCD630-like plasmids in these strains raises some interesting questions. There is little to no information on the function of these plasmids in *C. difficile* cells. The plasmids from the pCD630-family lack characterized determinants for antimicrobial resistance and are therefore unlikely to play a major role in drug resistance. Instead, they appear to harbor phage remnants or (partial) mobile genetic elements. It is documented that (pro)phages can modulate the expression of the major toxins [45, 46], affect the expression of cell wall proteins [47] and are up-regulated during infection [48]; a role in virulence of *C. difficile* is therefore certainly conceivable.

This study has only looked at plasmids of the pCD630 family and found that it occurs among diverse *C. difficile* strains. Based on our limited survey, we found plasmids in 5 different PCR ribotypes, and in strains of different toxinotypes (including both toxigenic and non-toxigenic strains). It will be of interest to see if the pCD630-family of plasmids is the most common, or that other plasmids are equally widely distributed. A broad survey of available genome sequences will likely reveal other families of plasmids and some of these may be limited to specific strains or clades of *C. difficile*.

The distribution of pCD630-like plasmids suggests that this family was acquired early during the evolution of *C. difficile*, or that the plasmid is capable of horizontal transfer. The pCD630-like plasmids do not encode any characterized conjugation proteins (Figure 2); however, they might be transferable dependent on other mobile elements or conjugative plasmids. Of note in this respect is that the mobile element ICE*Bs1* (which is related to Tn*916*, a conjugative transposon common in *C. difficile*) can mobilize plasmids [49], the pathogenicity locus of *C. difficile* can get transferred by a so far unidentified mechanism likely to rely on integrated conjugative elements [50] and in archaea vesicle-mediated plasmid transfer has been documented [51].

We found that pCD630-like plasmids are compatible with different replicons (Figure 1C). To our knowledge, no plasmid incompatibility has been described for *C. difficile* and sequence analysis did not reveal clear candidate genes for an incompatibility system in the plasmids analyzed. Considering the high plasmid prevalence (Figure 3), and the fact that existing genetic tools for *C. difficile* depend on the conjugative transfer of shuttle plasmids with a pCB102 or pCD6 replicon [5], one can wonder whether some strains are refractory to genetic manipulation due to the presence of plasmids from an incompatible plasmid group.

## Conclusions

In this study we showed that plasmid pCD630 from strain 630 is the paradigm of a family of plasmids that is defined by a module that encodes a conserved helicase. Most of the family members belong to a group that is larger than pCD630, and that differ in their accessory module. Plasmids from the pCD630-family are present in diverse *C. difficile* strains. Our data warrant further investigation of the role of pCD630-like plasmids – and plasmids in general - in *C. difficile* biology.

## Acknowledgements

The following people are acknowledged for helpful discussions: Y. Anvar, W. Meijer, M. Krupovic and the Experimental Bacteriology laboratory of E.J. Kuijper at the LUMC. We thank the Wellcome Trust Sanger Institute for sharing unpublished raw data that was used in this work via the European Nucleotide Archive. The European Nucleotide Archive support desk is acknowledged for updating the AM180356 record to reflect the findings of this study. This work was supported by the Netherlands Organisation for Scientific Research [VIDI 016.141.310] and Departmental Funds to W.K.S.

